# A Plug-and-Play Platform for Automated Azapeptide Synthesis: Case study of Azapeptide-based GLP-1

**DOI:** 10.1101/2025.03.03.641232

**Authors:** Mingzhu He, Kai Fan Cheng, Yousef Al-Abed

**Author notes:** Yousef Al-Abed, **E-mail:**. **Author Contributions:** M.H., K.F.C., and Y.A. designed the study; M.H., and K.F.C. ran experiments; M.H., and K.F.C. analyzed all data; and M.H. wrote the initial draft of the manuscript; and K.F.C., Y.A. contributed to manuscript revision and editing. **Competing Interest Statement:** K.F.C., Y.A. are on a patent application held by the Feinstein Institutes for Medical Research (FIMR) related to the azapeptide synthesis platform-Preparation of O-benzotriazole and O-imidazole synthons for use in the synthesis of peptidomimetics including azapeptides, WO2020227594 A1 2020-12-11 (active). Y.A. is inventor (FIMR) on Synthesis and uses of peptidomimetics including azapeptides, WO2020227588 A1 2020-11-12 (active). Y.A. is inventor (FIMR) on Aza GLP-1-based therapeutic analogues. United States, Provisional filing USSN 63/609,975. All the other authors declare no competing interests.

## Abstract

Azapeptide modification, achieved by substituting backbone α-carbons with nitrogen atoms to form enzyme-resistant semicarbazide bonds, can markedly enhance peptide stability and therapeutic potential. However, broad application has been constrained by two major synthetic challenges: the lack of suitable building blocks and the reduced nucleophilicity of the semicarbazide amino group, which limits post-coupling efficiency in automated solid-phase peptide synthesis (SPPS). Here, we describe a fully automated azapeptide synthesis platform that employs Fmoc-protected benzotriazole esters as bench-stable, pre-activated aza-amino acid building blocks. Microwave-assisted synthesis was integrated to accelerate aza-residue incorporation and improve coupling efficiency. This plug-and-play approach enables rapid solid-phase assembly of azapeptides, substantially reducing reaction times and improving yields. To demonstrate its utility, we synthesized azapeptide analogues of glucagon-like peptide-1 (GLP-1) with targeted substitutions at protease-sensitive sites, achieving enhanced stability as demonstrated in our recent study (bioRxiv 2025). This automated platform overcomes long-standing barriers in azapeptide chemistry, providing a scalable route for the rapid generation of stabilized peptide therapeutics.

**Significance Statement:** Azapeptides are a promising class of peptidomimetics with enhanced enzymatic stability and therapeutic potential. Yet, their synthesis has been limited by two persistent barriers: the lack of suitable, stable building blocks and the reduced coupling efficiency caused by the semicarbazide backbone. We developed a fully automated, microwave-assisted solid-phase synthesis platform that overcomes both challenges by using bench-stable benzotriazole ester aza-amino acid building blocks. This plug-and-play system enables efficient incorporation of aza-residues under standard SPPS conditions, reducing reaction times while improving yields and reproducibility. The resulting platform allows rapid generation of azapeptide libraries for biological screening and drug development, representing a major step toward scalable production of protease-resistant peptide therapeutics.

## Introduction

Azapeptides (1, 2) – peptides in which one or more backbone *α*-carbons are replaced by nitrogen atoms-represent a promising class of peptidomimetics that address the inherent instability of native peptides. This nitrogen substitution introduces a rigid trigonal geometry and enhances resistance to enzymatic hydrolysis. Despite their therapeutic potential, highlighted by the clinical success of Goserelin (3) (Zoladex®) for prostate and breast cancer and Atazanavir (4) (Reyataz®) for HIV infection, the broader development of azapeptides has been constrained by long-standing synthetic challenges. Traditional methods (2, 5, 6) typically rely on the activation of hydrazine derivatives with carbonyl-donating reagents, such as phosgene or chloroformates, which are often unstable, and cumbersome to process (Figure 1a). Alternative strategies, including submonomer azapeptide synthesis (7, 8) and customized solid-phase peptide synthesis (SPPS) (2), have advanced the field but often yield poor coupling efficiency and low overall recovery under harsh reaction conditions. The two main obstacles remain the lack of stable, pre-activated aza-amino acid building blocks and inefficient post-coupling reaction due to the reduced nucleophilicity of the terminal semicarbazide nitrogen.

**Figure 1.**
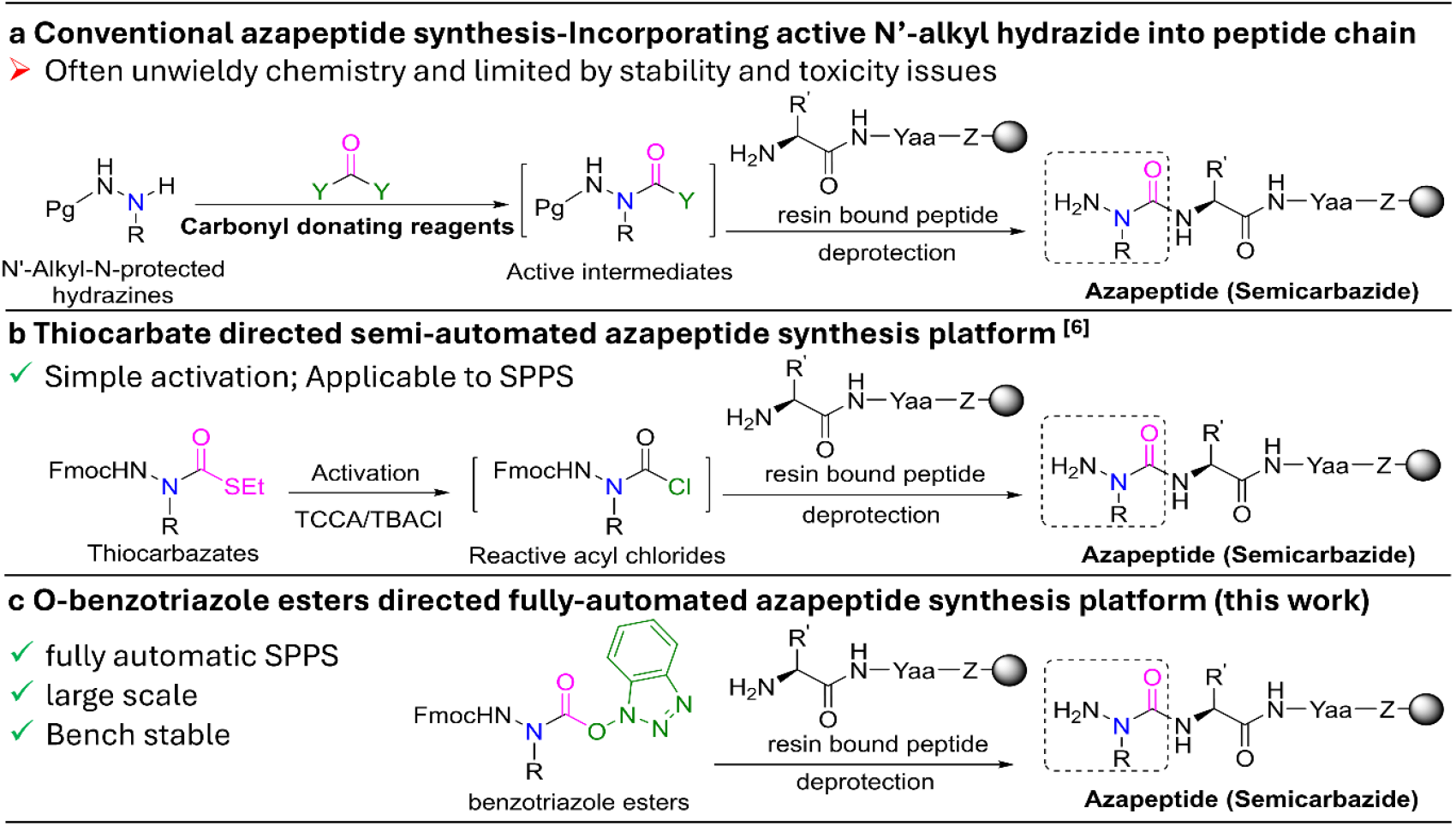
Current azapeptide synthetic strategies. a. conventional azapeptide synthesis - activation of N’-alkylated hydrazine derivatives with carbonyl donating reagents (such as chloroformates, p-nitrophenyl chloroformate, bis(2,4-dinitrophenyl) carbonate, carbonyldiimidazole (CDI), 1,1′-Carbonyl-di-(1,2,4-triazole) (CDT), bis(pentafluorophenyl) carbonate, and N,N’-disuccinimidyl carbonate (DSC); b. thiocarbazate-based azapeptide synthesis - simple activation (5-10 mins at r.t.) of shelf-stable thiocarbazate building block (9), compatible with solid phase chemistry; c. O-benzotriazole esters-based azapeptide synthesis - bench stable, pre-activated aza precursors, providing fully automation strategy in solid phase azapeptide synthesis (21). Abbreviations – Pg: Boc, Ddz, Phth and Fmoc; R, R’: H, amino acids side chain; Yaa: amino acids; Z: O (Wang resin), NH (Rink amide resin)

In our previous work, we introduced a semi-automated synthetic platform based on thiocarbazate building blocks as stable, activatable precursors, enabling the generation of azapeptide libraries (9) (Figure 1b) with enhanced stability and biological activity, as demonstrated in model peptides, FSSE and bradykinin. Here, we report a fully automated solid-phase azapeptide synthesis platform (Figure 1c) that overcomes both major synthetic barriers. The method employs Fmoc-protected benzotriazole esters as bench stable, pre-activated aza-amino acid building blocks compatible with standard SPPS automation and microwave-assisted coupling. This plug-and play system accelerates azapeptide assembly, reduce by-products and improve yields. To demonstrate its utility, we applied the platform to strategically edit the dipeptidyl peptidase-4 (DPP-IV) cleavage site in glucagon-like peptide-1 (GLP-1) (10), generating four aza-substituted GLP-1 analogues (11).

## Results and Discussion

### Synthesis of aza-benzotriazole ester building blocks

Unlike natural amino acids, aza-amino acids are inherently unstable, requiring alternative activation strategies (5, 12). Hydroxybenzotriazoles, such as 1-hydroxybenzotriazole (HOBt), generate reactive ester intermediates in situ during peptide synthesis (13, 14). We hypothesized that HOBt could react with aza-amino acyl chloride to form a stable carbazates suitable for isolation, storage, and subsequent coupling without further activation. Sixteen Fmoc-azaAA-benzotriazole ester (Fmoc-azaAA-OBt) building blocks (**2a-2p**) were synthesized by treating N-Fmoc-N’-substituted hydrazines (**1a-1p**) (5, 6) with 1.5 equivalents of phosgene to form transient acyl chlorides, immediately immediately followed by trapping with 1.5 equivalents of HOBt in DMF. Reactions proceeded efficiently, yielding bench-stable building block in 73.3-96.1%, Table S1, Supporting Information). X-ray crystallographic of compound Fmoc-azaVal-OBt (**2j**) (Supplementary Note; CCDC-2486574) confirmed the structural representative of this class. These building blocks maintained long-term stability in the solid state at -20 °C and retained >95% integrity in DMF at room temperature for 24 h. Notable exceptions included glycine (azaglycine carbazate undergoes intramolecular cyclization to form oxadiazalone (15)), serine, threonine, and cysteine (inherently incompatible with azapeptide chemistry, therefore, not investigated). These results demonstrate that aza-benzotrizole esters overcome the key stability limitation of conventional aza-amino acids.

### Automated incorporation of aza-amino acids via standard SPPS

To evaluate reactivity, aza-benzotrizole esters were incorporated into tyrosine pre-loaded Wang resin as a model system. Using standard Fmoc/tBu SPPS protocols, 5 equivalents of Fmoc-azaAA-OBt (**2a-p**) were coupled to the resin at room temperature for 16h. Over 80% incorporation occurred at 1 h, with optimal conversion after 16 h. Kaiser tests were used to monitor reaction progress. All sixteen benzotriazole esters were successfully coupled, yielding azadipeptides (**4a-4p**) with 85.4%-100% conversion and 62.7-95.1% crude purity (Table 1). The resulting azadipeptides were characterized by HPLC, HRMS and NMR analyses (see supporting information, section 2.3). Notably, azaWY-OH (**4f**) and azaYY-OH (**4o**) side chains remain sensitive (6, 9) and required optimized TFA cleavage conditions (Table 1, supporting information, section 2.3). In sum, aza-benzotriazole esters uniquely combine bench stability with high reactivity, enabling automated SPPS of azadipeptides with high reliability.

**Table 1.**
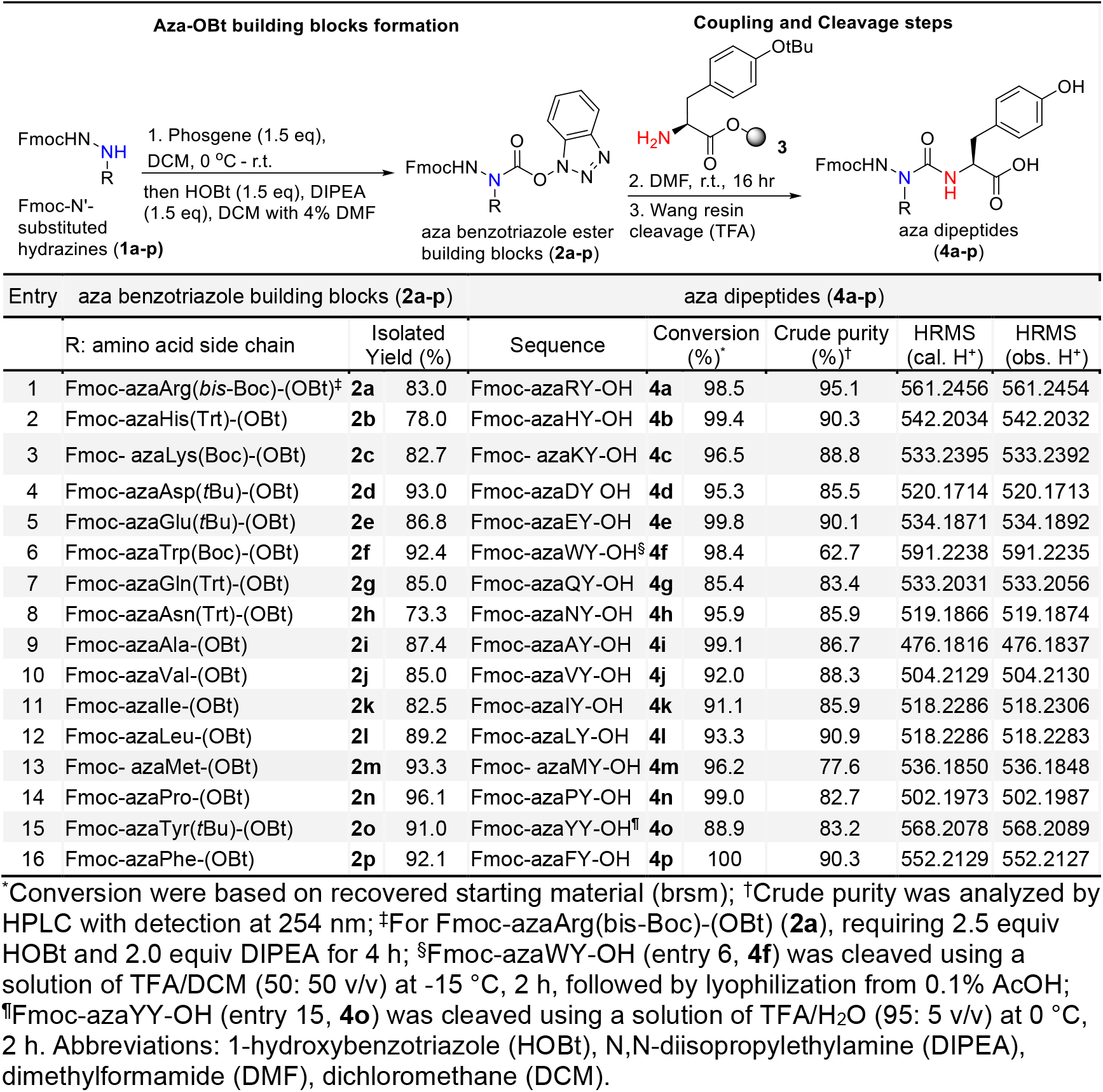
Automated incorporation of aza-moieties: synthesis of azadipeptides using 16 benzotriazole ester building blocks via standard SPPS. Reagents and conditions: (1) Synthesis of benzotriazole esters building blocks: Phosgene (1.5 equiv), DCM, 0 °C – r.t., 0.5 h then HOBt (1.5 equiv), DIPEA (1.5 equiv), 4% DMF in DCM, r.t., 0.5 h; (2) Integration of the aza-amino acids in SPPS: NH_2_-Tyr(tBu)-Wang Resin (1 equiv), aza-amino acid OBt (5 equiv, 0.2 M), DMF, r.t., 16 h; (3) Cleavage step: TFA/H_2_O/TIPS (95:2.5:2.5. v/v/v), r.t., 2 h, 10-25 ml/g of resin. Synthesis and characterization of these aza-amino acid building blocks and azadipeptides are detailed in Supporting Information

### Optimization of post-coupling conditions to minimize incomplete acylation in automated azapeptide synthesis

Efficient chain elongation after aza-residue incorporation remains one of the most persistent barriers in azapeptide chemistry. The semicarbazide nitrogen of an aza-amino acid is a weaker nucleophile than the α-amino group of natural amino acids, often resulting in incomplete coupling during the solid-phase synthesis (5, 16). Overcoming these challenges is essential for enabling full automation and reproducibility of azapeptide assembly.

Following Fmoc deprotection, semicarbazide moieties (Figure 1) were acylated to extend peptide sequences (Table 2) (5, 16). Using azaF^2^I dipeptide (**6**) as a model aza-amino acid residue, we assessed its coupling performance with natural amino acids of diverse side chains, employing multiple standard coupling reagents (Table 2a, entry 9-14). HCTU-mediated coupling achieved 76.3–91.5% conversion for flexible residues, with isolated yields of 50.0– 79.5%, comparable to those of natural tripeptides (supplementary Table S2). However, sterically hindered residues such as valine and isoleucine exhibited low coupling efficiencies, with conversions ranging from 17.4% to 35.4% (Table 2a, entry 8 and 9; supplementary Table S3). Alternative coupling reagents, such as DIC, HDMA, T3P®, COMU, improved outcomes only marginally, requiring multiple cycles and prolonged reaction times (Table 2a, entry 10-13). Although various groups (17) have made progress in overcoming these obstacles using halogenating reagents, these methods are not compatible with automated SPPS. These findings underscore the persistent synthetic challenges posed by steric hindrance during aza-amino acid acylation and the need for SPPS-compatible solutions.

**Table 2.** Automated acylation of the resin-linked azapeptides: model reactions of coupling natural amino acid to aza-amino acid under traditional and microwave-assisted SPPS conditions. Reagents and conditions: a. Synthesis of azaPhe tripeptide **8a-i**.1) Coupling step, NH_2_-Ile-Wang Resin (**5**, 1 equiv), Fmoc-azaF-OBt (**2p**, 5 equiv), DMF, r.t., 16 h; 2) Fmoc removal, 20% piperidine solution in DMF, 2.5 min; 3) Acylation step (entry 1-13), Fmoc-amino acids **7a-i** (5 equiv), coupling reagents (10 equiv) with 12 equiv of NMM in DMF under traditional solid phase peptide synthesizer; entry 14, Fmoc-L-Ile-OH (**7i**, 5 equiv), DIC (10 equiv)/ OxymaPure® (5 equiv) in DMF using microwave assisted peptide synthesizer. 4) Fmoc removal, 20% piperidine solution in DMF, 2.5 min; b. Synthesis of azaF-decapeptide **12** using optimized conditions. Traditional peptide synthesizer: 1) Coupling step, NH_2_-IAKVVEGG-Rink amide Resin (**9**, 1 equiv), Fmoc-azaF-OBt (**2p**, 5 equiv), DMF, r.t., 16 h; 2) Fmoc removal, 20% piperidine solution in DMF, 2.5 min; 3) Acylation step, Fmoc-L-Glu (OtBu)-OH **11** (5 equiv), COMU (10 equiv) with 12 equiv of NMM in DMF, 4 coupling cycles, 8 h; Microwave Peptide Synthesizer: 1) Coupling step, NH_2_-IAKVVEGG-Rink amide Resin (**9**, 1 equiv), Fmoc-azaF-OBt (**2p**, 5 equiv), DMF, 60 °C MW, 1 h or 90 °C MW, 4 min; 2) Fmoc removal, 10% piperidine solution in DMF, 1 min at 90 °C MW.; 3) Acylation step, Fmoc-L-Glu (OtBu) **11** (5 equiv), DIC (10 eq) /OxymaPure® (5 eq) in DMF, 2 coupling cycles, 2 h at 60 °C MW. Synthesis and characterization of these azatripeptides are detailed in Supporting Information.

| Entry | azaPhe-tripeptides | R (natural amino acid residues) | Coupling reagent (10 eq) | Rxn time/Temp. | Conversion (%)<sup>*</sup> | Crude purity (%)<sup>†</sup> | Crude yields (%) | Isolated yields (%) | Purity (%)<sup>†</sup> |
| --- | --- | --- | --- | --- | --- | --- | --- | --- | --- |
| 1 | HazaF<sup>2</sup>I-OH (**8a**) | Histidine | HCTU | 2 h / coupling cycle<br>2 cycles, 4 h, r.t. | 90.0 | 72.7<sup>‡</sup><br>(3:1) | 83.1 | 50.0 | 99.1<sup>‡</sup><br>(3:1) |
| 2 | RazaF<sup>2</sup>I-OH (**8b**) | Arginine | HCTU | 2 h / coupling cycle<br>5 cycles, 10 h, r.t. | 76.3 | 76.7 | 70.5 | 62.1 | 98.4 |
| 3 | YazaF<sup>2</sup>I-OH (**8c**) | Tyrosine | HCTU | 2 h / coupling cycle<br>4 cycles, 8 h, r.t. | 88.1 | 76.3 | 83.2 | 79.5 | 90.3 |
| 4 | WazaF<sup>2</sup>I-OH (**8d**) | Tryptophan | HCTU | 2 h / coupling cycle<br>3 cycles, 6 h, r.t. | 82.7 | 80.6 | 77.8 | 53.6 | 98.6 |
| 5 | PazaF<sup>2</sup>I-OH (**8e**) | Proline | HCTU | 2 h / coupling cycle<br>4 cycles, 8 h, r.t. | 87.1 | 88.7 | 96.7 | 69.1 | 97.6 |
| 6 | MazaF<sup>2</sup>I-OH (**8f**) | Methionine | HCTU | 2 h / coupling cycle<br>5 cycles, 10 h, r.t. | 86.7 | 67.0 | 78.4 | 61.0 | 96.1<sup>‡</sup><br>(12:1) |
| 7 | AazaF<sup>2</sup>I-OH (**8g**) | Alanine | HCTU | 2 h / coupling cycle<br>3 cycles, 6 h, r.t. | 91.5 | 70.5 | 90.9 | 77.4 | 97.0 |
| 8 | VazaF<sup>2</sup>I-OH (**8h**) | Valine | HCTU | 2 h / coupling cycle<br>4 cycles, 8 h, r.t. | 17.4 | N.A. | N.A. | N.A. | N.A. |
| 9 | IazaF<sup>2</sup>I-OH (**8i**) | Isoleucine | HCTU | 2 h / coupling cycle<br>4 cycles, 8 h, r.t. | 35.4 | N.A. | N.A. | N.A. | N.A. |
| 10 |  |  | DIC |  | 18.0 | N.A. | N.A. | N.A. | N.A. |
| 11 |  |  | T3P® |  | 28.3 | N.A. | N.A. | N.A. | N.A. |
| 12 |  |  | HDMA |  | 44.3 | N.A. | N.A. | N.A. | N.A. |
| 13 |  |  | COMU |  | **68.8** | 66.0 | 75.8 | **40.8** | 95.0 |
| 14 |  |  | DIC/Oxyma Pure®<br>(10 eq/5 eq) | 1 h /coupling cycle<br>4 cycles, 4 h,<br>60 °C MW. | **88.3** | 61.2 | 90.8 | **63.0** | 98.7 |

| SPPS | Rxn temperature | azaPhe-semicarbazide (**10**) |  |  |  | azaPhe-azadecapeptide (**12**) |  |  |  |
| --- | --- | --- | --- | --- | --- | --- | --- | --- | --- |
|  |  | Rxn time | Conversion (%)<sup>*</sup> | Crude purity (%)<sup>§</sup> | Coupling reagent | Rxn time | Conversion (%)<sup>*</sup> | Crude purity (%)<sup>†</sup> | Side product (%) |
| Traditional Peptide Synthesizer | r.t. (25 °C) | 16 h | 99.7 | 93.4 | COMU (10 eq) | 2 h / coupling cycle<br>4 cycles, 8 h, r.t. | 85.6 | 69.3 | 7.5 |
| Microwave Peptide Synthesizer | 60 °C MW. | 1 h | 99.9 | 94.7 | DIC/Oxyma Pure®<br>(10 eq/5 eq) | 1 h / coupling cycle<br>2 cycles, 2 h | 95.9 | 83.8 | No |
|  | 90 °C MW. | 4 min | 99.3 | 88.4 | N.A. | N.A. | N.A. | N.A. | N.A. |

<sup>\*</sup> Conversion was based on recovered starting material (brsm); <sup>†</sup> Crude and final purity were analyzed by HPLC with detection at 215 nm; <sup>‡</sup> Total purity includes peptide isomers. Relative isomer ratios are indicated in parentheses. <sup>§</sup> Crude purity of **10** were analyzed by HPLC with detection at 215 nm as Fmoc protected semicarbazide; Abbreviations: (2-(6-chloro-1-H-benzotriazol-1-yl)-1,1,3,3-tetramethylaminium) hexafluorophosphate (HCTU), N,N-

To overcome these obstacles, we investigated the impact of elevated temperature on post-aza coupling efficiency during isoleucine acylation (Table 2a, entry 14). Microwave-assisted acylation at 60 °C using DIC/OxymaPure over four cycles significantly enhanced performance, achieving 88.3% conversion (a 20% increase compared to COMU), a 63.0% isolated yield, and halving the reaction time to 4 hours. This optimization effectively mitigates steric hindrance and incomplete acylation, providing a robust and scalable strategy for automated azapeptide synthesis.

### Fully automated synthesis of an azadecapeptide and aza-GLP-1 analogues

Based on these findings, we developed a fully automated microwave-assisted SPPS workflow for longer azapeptides: natural amino acids were coupled at 90 °C (2 min, DIC/Oxyma), while aza-residues were incorporated at 60 °C (60 min), and semicarbazide acylation cycles were adjusted based on steric hindrance of subsequent amino acid. To validate the protocol, a 10-amino acid GLP-1 fragment (18) was synthesized. As shown in Table 2b, microwave irradiation reduced the azaPhe incorporation time from 16 h at room temperature to 1 h at 60 °C while maintaining >99% conversion and increasing crude purity of semicarbazide (**10**) to 94.7%. Similarly, the acylation time of Fmoc-Glu-OH (**11**) was reduced from 8 h to 2 h, and the crude purity of the resulting azadecapeptide (**12**) improved from 69.3% to 83.8%.

To further investigate the limits of acceleration, we conducted preliminary studies on aza-amino acid incorporation at 90 °C—the standard coupling temperature for natural amino acids. Remarkably, initial results (Table 2b) showed complete incorporation of azaPhe in just 4 minutes, with 99.3% conversion and 88.4% crude purity of azaF^2^ nonapeptide **10**, and no detectable degradation of the azaPhe-benzotriazole ester or formation of side products. This result represents a substantial improvement over the 60 °C protocol. In addition, elevated temperatures such as 90 °C may further enhance acylation efficiency of sterically hindered residues. While preliminary, these findings underscore the potential of our platform to enable rapid and efficient azapeptide synthesis.

Encouraged by the successful synthesis of the azadecapeptide GLP-1 fragment, we applied our platform and methods to construct GLP-1 azapeptide analogues. GLP-1, a 30-or 31-residue hormone that enhances glucose-dependent insulin secretion, suppresses glucagon release and slow gastric emptying. Its therapeutic relevance in type 2 diabetes and obesity has led to several FDA-approved analogues (10), including semaglutide (19), which incorporates two key modifications: replacing alanine at position 8 with alpha-aminoisobutyric acid (Aib), an unnatural amino acid, to prevent dipeptidyl peptidase-4 (DPP-IV) degradation and introducing a lipid chain to lysine at position 26 to facilitate albumin binding and slow release (Table 3a). Recently, Kumar et al. (20) demonstrated that replacing key residues with aza-amino acids stabilizes GLP-1 (7-36) amide, semaglutide and gastric inhibitory polypeptide (GIP) analogues by preventing DPP-IV-catalyzed hydrolysis and inactivation. Although these results were encouraging, the reported synthetic methods utilized cumbersome protocols – including multiple carbonyl donors such as triphosgene, disuccinimidyl carbonate (DSC) and 4-nitrophenyl chloroformate, requiring prolong reaction times, non-SPPS-compatible conditions (-10°C), and labor-intensive steps.

**Table 3.**
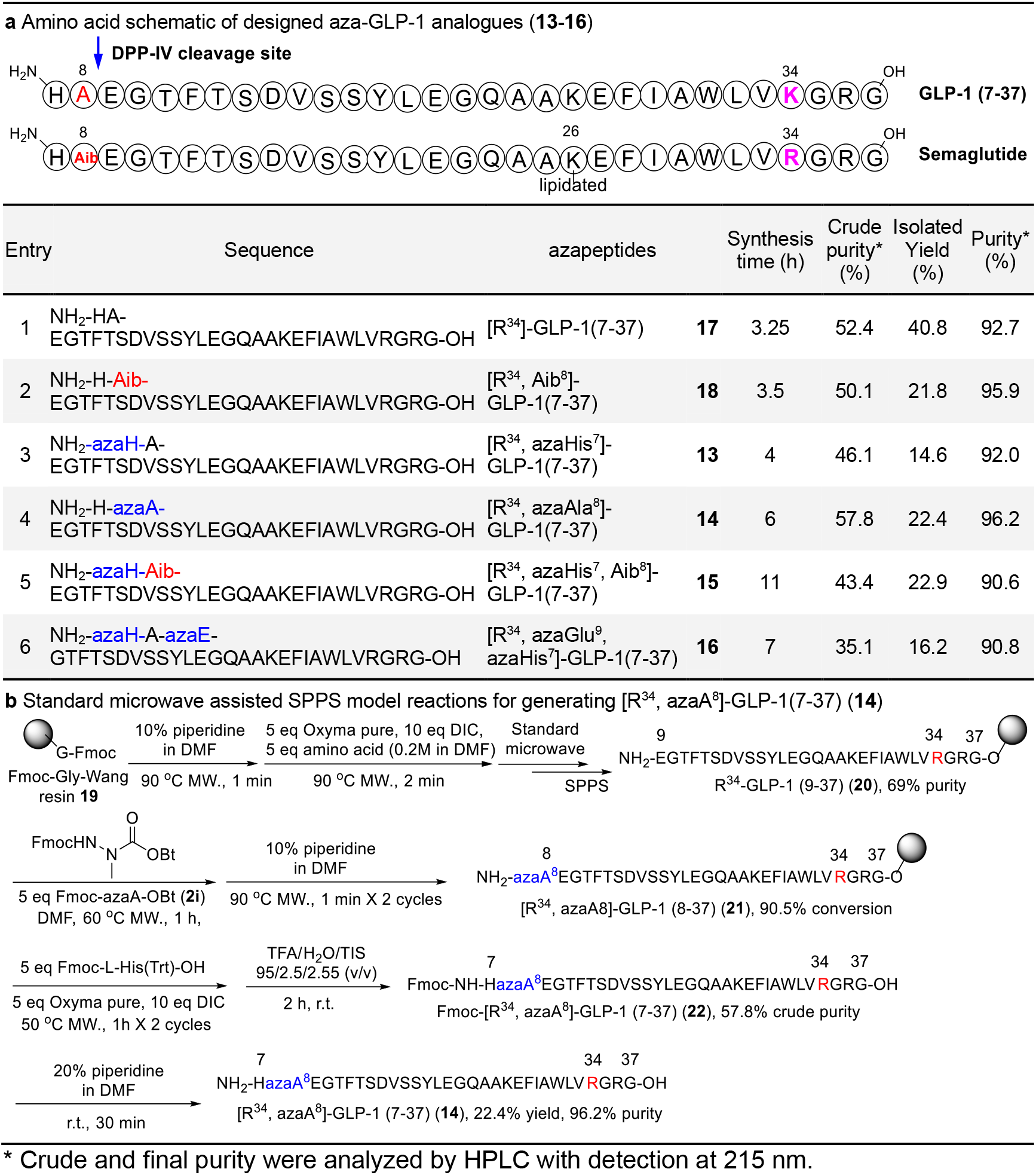
Fully automatic synthesis of aza-Glucagon-like peptide-1 (GLP-1) analogues: trial application of optimized azapeptide synthesis platform. a. Amino acid schematic of designed aza-GLP-1 analogues showing aza-amino acid substitutions around DPP-IV cleavage sites; summary of GLP-1 and GLP-1-based azapeptide analogues (**13-16**); b. Standard microwave assisted SPPS for generating [R^34^, azaA^8^]-GLP-1(7-37) (**14**) was presented as model reactions. Synthesis and HPLC/HRMS of aza-Glucagon-like peptide-1 (GLP-1) analogues are detailed in Supplementary Information (section 2.6).

In contrast, using our fully automated platform, we successfully synthesized four R^34^ aza-GLP-1 (7–37) analogues via microwave-assisted standard solid-phase peptide synthesis, incorporating aza-alanine, aza-histidine, or aza-glutamic acid near the DPP-IV cleavage site. The crude purities ranged from 35.1% to 57.8%, with isolated yields between 14.6% and 22.9% (Table 3a, **13**–**16**). [R^34^, azaAla^8^]-GLP-1(7–37) (**14**) was selected as a representative analogue, with its synthetic scheme illustrated in Table 3b. This analogue was obtained in 22.4% yield within 6 hours—comparable to the Aib^8^-GLP-1 analogue (**18**), the unlipidated semaglutide, which was synthesized in 3.5 hours with a 21.8% yield. In general, single aza-modified GLP-1 analogues were synthesized in 4–6 hours, while multi-aza analogues required 7–11 hours, representing a substantial reduction in total synthesis time compared to previously reported methods. Overall, our results establish a scalable and reproducible automated platform broadly applicable to the synthesis and optimization of long-sequence therapeutic peptides.

To conclude, we have developed a novel fully automatic solid phase synthetic platform for azapeptides synthesis utilizing aza-benzotriazole ester building blocks. These results highlight the robustness and reliability of our platform for synthesizing challenging peptides like GLP-1. The availability of on-demand syntheses of peptides resistant to proteolytic degradation enzymes marks a significant advancement in the pharmaceutical industry. Our approach has the potential to enable the more rapid synthesis and future development of novel azapeptide-based therapeutics, which can address unmet medical needs. Preliminary high-temperature incorporation studies suggest that further acceleration of synthesis is feasible, setting the stage for even more efficient azapeptide library generation in future work.

## Materials and Methods

### General procedure for the synthesis of benzotriazole amino acid building blocks

To a solution of substituted hydrazine (2 mmol, 1 eq, 0.08 M) in anhydrous DCM (25 mL), phosgene (3 mmol, 1.5 eq, 15% solution in toluene) was added at 0 °C. The reaction mixture gradually warmed up to room temperature and stirred for 0.5 hour (monitoring by TLC). Then, the reaction mixture was stopped by evaporating the excess volatiles and the crude material was dried under vacuum pump for 1 hour to give the corresponding acyl chloride which was re-dissolved in anhydrous DCM (20 mL). To the above solution, a HOBt 0.5 M solution (3 mmol, 1.5 eq in 1.0 mL DMF and 5.0 mL DCM) and DIPEA (3 mmol, 1.5 eq) were added. The reaction mixture was stirred at room temperature for 0.5 hour (monitoring by TLC), then was quenched with water (25 mL) and extracted with EtOAc (25 mL x 4). The combined organic layer was washed with brine (25 mL), dried over Na_2_SO_4_, filtered and evaporated under vacuum. The crude residue was purified by CombiFlash® Teledyne Isco Chromatography with a linear gradient from hexanes: EtOAc [95:5] to hexanes: EtOAc [5:95] in 20 minutes. Characterization of 16 benzotriazole esters can be found in Supporting Information.

### General procedure for solid phase peptide synthesis (SPPS) at room temperature

All solid phase peptide couplings were performed at room temperature using Tribute® Peptide synthesizer from Gyros Protein Technologies, Inc following standard protocol which is described sequentially below:

1. Swelling: The resin (Rink amide or preloaded Wang resin) was swelled three time successively for 10 min each time in DMF.
2. Fmoc Cleavage: the protected amino acid/peptide or azapeptide was shaken for 2.5 min with 20% piperidine solution in DMF to remove the Fmoc group. The process was repeated twice, followed by several washing steps with DMF (5 times).
3. Amino acid coupling: 5 equivalents of the next acylating component (Fmoc-protected amino acid), 5 equivalents of HCTU (coupling reagent), and 10 equivalents of N-methylmorpholine (base) were used to add the next amino acid in the sequence. This step is fully automated and was run according to the software installed on the Tribute® synthesizer. The amino acid including coupling reagent was delivered to the reaction vessel from the specified loading position upon dissolution. Then the base was added as 0.4 M solution in DMF, the total volume of solvent was adjusted to give 0.2 M solution. The coupling time was set to 10 min shaking followed by drainage then washing steps. Step 2 and step 3 were repeated until the desired peptide sequence was achieved.
4. Washing: repeated washing steps were performed after each cleavage or coupling event using DMF as solvent (5 times). At the final coupling or cleavage steps additional washing with DCM was performed (5 times) to remove any trace of DMF. The process was usually followed by a drying step.
5. Cleavage from the Resin: 3.0 mL of a freshly made solution of TFA/H_2_O/TIPS (95:2.5:2.5. v/v/v) was added at 25 °C to a 0.1 mmol of resin (10-25 mL/g of resin). The mixture was shaken for 2 h at 25 °C, filtered, and the remaining resin was further washed with a 0.5-1.0 mL of TFA solution. The excess TFA solution was removed under nitrogen gas. The filtrate was precipitated by adding 30 mL of ice-cold ether. Upon centrifugation, the resulting solid was dissolved in 4 mL of 1:4 solution of CH_3_CN: H_2_O. The resulting solution was lyophilized.
6. Purification: purification of the peptidomimetics were performed on a preparative HPLC purification system (Waters Prep 150 LC system combining 2545 Binary Gradient Module using XSelect Peptide CSH C18 OBD Prep Column, 130Å, 5 µm, 19 mm X 150 mm. Chromatography was performed at ambient temperature with a flow rate of 30 mL/min with a linear gradient from water (0.05% TFA): CH_3_CN (0.05% TFA)[95:5] to water (0.5% TFA): CH_3_CN (0.05% TFA) [5:95] in 18minutes, monitored by 2998 Photodiode Array (PDA) Detector UV at 254 nm and/or 215 nM.

### General procedure for solid phase peptide synthesis (SPPS) at 90 ^°^C with microwave heating

All solid phase peptide couplings were performed at 90 °C using Liberty Blue 2.0 Microwave Peptide synthesizer from CEM Corporation following standard protocol which is described sequentially below:

1. Swelling: The resin (Rink amide or preloaded Wang resin) was swelled for 5 min in DMF at room temperature.
2. Fmoc Cleavage: the protected amino acid/or peptide was mixed for 1 min with 10% piperidine solution in DMF at 90 °C to remove the Fmoc group. Followed by several washing steps with DMF (4 times).
3. Amino acid coupling: 5 equivalents of the next acylating component (Fmoc-protected amino acid), 5 equivalents of OxymaPure® (coupling reagent), and 10 equivalents of DIC (coupling reagent) were used to add the next amino acid in the sequence. This step is fully automated and was run according to the software installed on the synthesizer. The coupling time was set to 2 min mixing at 90 °C followed by drainage step. Step 2 and step 3 were repeated until the desired peptide sequence was achieved.
4. Cleavage and purification were performed as described in Supporting Information 2.2.1 (step 5 and 6)

### General procedure for automated integration of the aza-amino acid in SPPS

Automated incorporation of the aza-amino acid moieties in solid phase was performed using two different protocols; the Kaiser test can be used during synthesis to monitor coupling and adjust the reaction time as needed:

1. Integration of the aza-amino acid in SPPS at room temperature using Tribute® Peptide synthesizer. Wang or Rink amide amino acid-loaded resin (0.1 mmol) was swelled for 10 min in DMF (2.5 mLx3), followed by Fmoc cleavage using 20% solution of piperidine in DMF (2.5 mL x 2, 2.5 min per cleavage cycle). After successive washes with DMF (2.5 mL x 5, 30 seconds per wash), Fmoc-aza-OBt building block (0.5 mmol, 5 equiv, 0.2M) in DMF (2.5 mL) was automatically transferred into the reaction vessel. The reaction mixture was shaken for 16 h at 25 °C, followed by 5 cycles of washing and then drying. A small amount of the resin was cleaved using a freshly made solution of TFA/H_2_O/TIPS (95:2.5:2.5. v/v/v) and the resulting azadipeptide was analyzed by HPLC.
2. Integration of the aza-amino acid in SPPS at elevated temperature using CEM Microwave Peptide Synthesizer. Wang or Rink amide amino acid-loaded resin (0.1 mmol) was swelled for 10 min in DMF (10 mL), followed by Fmoc cleavage using 10% piperidine solution in DMF at 90 °C to remove the Fmoc group. Followed by washing steps with DMF (4.0 mL x 4), Fmoc-aza-OBt building block (0.5 mmol, 5 equiv, 0.2 M) in DMF (2.5 mL) was automatically transferred into the reaction vessel. The coupling time was set to 30-60 min mixing at 60 °C followed by drainage step. A small amount of the resin was cleaved using a freshly made solution of TFA/H_2_O/TIPS (95:2.5:2.5. v/v/v) and the resulting azadipeptide was analyzed by HPLC.

### General procedure for the automated acylation of resin-linked azapeptide in SPPS

Functionalization of the aza-amino acid moieties of the solid phase was performed using two different protocols:

1. Coupling to the aza-amino acid on the peptidyl chain with HCTU or other coupling reagents including DIC, T3P®, HDMA and COMU was performed at room temperature using Tribute® Peptide synthesizer. The Fmoc protected aza-amino acid bound to the peptidyl chain (based on 0.1 mmol peptide) was treated with 20% piperidine solution in DMF (2.5 mL), the suspension was shaken for 2.5 min, and then the de-protection solution was drained. The Fmoc-cleavage protocol was repeated three times to ensure complete de-protection. After removing Fmoc group from the peptidyl chain, the resin was suspended in 0.4 M solution of NMM in DMF (3 mL, 12 eq, 1.2 mmol) with designated amino acid (5 eq, 0.5 mmol) and coupling reagent (10 eq, 1.0 mmol). The resulting suspension was shaken for 2 h at 25 °C, followed by drainage and washes with DMF (2.5 mL x 3) and DCM (2.5 mL x 5). The cycle was repeated multiple times based on the conversion rate, which was established by HPLC analysis after a cleavage of small amount of the resin.
2. Coupling to the aza-amino acid on the peptidyl chain with OxymaPure/DIC was performed at elevated temperature using CEM Microwave Peptide Synthesizer. The Fmoc protected aza-amino acid bound to the peptidyl chain (based on 0.1 mmol peptide) was mixed with 10% piperidine solution in DMF (4.0 mL) at 90 °C to remove the Fmoc group. Followed by several washing steps with DMF (4 mL x 4). The Fmoc-cleavage protocol was repeated twice to ensure complete de-protection. After cleavage of the Fmoc group from the peptidyl chain, the resin (0.1 mmol) was suspended in DMF (4.0 mL) with designated amino acid (5 eq, 0.5 mmol), OxymaPure (5 eq, 0.5 mmol) and DIC (10 eq, 1.0 mmol). The resulting suspension was mixed for 1 h at 60 °C, followed by drainage and washes with DMF. The coupling cycle was repeated multiple times based on the conversion rate, which was established by HPLC analysis after a cleavage of small amount of the resin.

### General procedure for the synthesis of GLP-1 azapeptide analogues

GLP-1 azapeptide analogues were synthesized on a liberty blue 2.0 microwave peptide synthesizer in fully automated mode. Azapeptide analogues (**13-18**) was synthesized from 0.1 mmol of Preloaded Fmoc-L-Gly Wang resin (100-200 mesh, 0.361 mmol/g) following the general procedure for solid phase peptide synthesis describe above. After standard SPPS, washes, and resin cleavage, GLP-1 azapeptide analogues (**13-18**) were analyzed by HPLC to confirm the crude yield and crude purity. Then the resulting crude material was purified using preparative C18 reverse-phase chromatography with a linear gradient. Following lyophilization, the isolated yield and final purity of each analogue were confirmed by analytical HPLC. **13-18** were collected and submitted to HRMS analysis. Synthesis and characterization of azapeptide analogues can be found in Supporting Information.

## Acknowledgements

We would like to thank Dr. Matthew Devany, Director of the NMR facility in the Department of Chemistry at Hunter College, for collecting NMR data and Dr. Barney Yoo from the Mass Spectrometry facility at Hunter College for acquiring HRMS data. The Single Crystal X-Ray Structure Determination has been performed at the X-Ray Crystallography facility in the Department of Chemistry at Hunter College, we thank Dr. Michelle C. Neary for her help. DoD funding for the X-ray machine at Hunter College is supported by the Air Force Office of Scientific Research under award number FA9550-20-1-0158. We thank Dr. Tom Coleman and Dr. John E. Moses for critical reading of the manuscript. We would like to thank Dr. Ahmad S. Altiti and Dr. Dharmendra S. Vishwakarma for helpful discussion and comments. This work was supported by funding from the Feinstein Institutes for Medical Research. Some of this work was supported by Northwell Health’s 2019 Innovation Challenge prize.

## Data Availability Statement

All relevant data to the manuscript generated for these studies are included in the article or supplemental information. The X-ray crystallographic coordinates for structures reported in this study have been deposited in the Cambridge Crystallographic Data Centre (CCDC), under deposition numbers CCDC-2486574.

**This preprint has not been peer-reviewed. All relevant raw data and supplemental information are available from the author upon request**.

## Notes

### Summary of Updates

This revision has been submitted to (1) update the copyright license to CC BY-NC-ND (Attribution-NonCommercial-NoDerivatives), (2) revise the title, Abstract, and Significance Statement for improved clarity (3) explicitly state that this preprint has not been peer-reviewed. Minor formatting adjustments and text edits have been made to enhance conciseness; however, no changes have been made to the figures or data presented in the original version.

## References

1. C. Proulx et al., Azapeptides and their therapeutic potential. Future Med. Chem. 3, 1139–1164 (2011).

2. K. F. Cheng, S. VanPatten, M. Z. He, Y. Al-Abed, Azapeptides -A History of Synthetic Milestones and Key Examples. Curr. Med. Chem. 29, 6336–6358 (2022).

3. C. J. Tyrrell et al., A Multicenter Randomized Trial Comparing the Luteinizing Hormone-Releasing Hormone Analogue Goserelin Acetate Alone and with Flutamide in the Treatment of Advanced Prostate Cancer. The Journal of Urology 146, 1321–1326 (1991).

4. P. J. Piliero, Atazanavir: a novel HIV-1 protease inhibitor. Expert Opinion on Investigational Drugs 11, 1295–1301 (2002).

5. A. Ploom, A. Mastitski, M. Arujoe, A. Troska, J. Jarv, Aza-peptides: expectations and reality. Proceedings of the Estonian Academy of Sciences 71, 241–254 (2022).

6. D. Boeglin, W. D. Lubell, Aza-Amino Acid Scanning of Secondary Structure Suited for Solid-Phase Peptide Synthesis with Fmoc Chemistry and Aza-Amino Acids with Heteroatomic Side Chains. J. Comb. Chem. 7, 864–878 (2005).

7. R. Chingle, C. Proulx, W. D. Lubell, Azapeptide Synthesis Methods for Expanding Side-Chain Diversity for Biomedical Applications. Acc. Chem. Res. 50, 1541–1556 (2017).

8. M. Bowles, C. Proulx, Solid phase submonomer azapeptide synthesis. Methods Enzymol. 656, 169–190 (2021).

9. A. Altiti et al., Thiocarbazate building blocks enable the construction of azapeptides for rapid development of therapeutic candidates. Nat Commun 13, 7127 (2022).

10. D. N. McBrayer, Y. Tal-Gan, Recent Advances in GLP-1 Receptor Agonists for Use in Diabetes Mellitus. Drug Dev. Res. 78, 292–299 (2017).

11. M. He et al., Azapeptide GLP-1 receptor agonist resists proteolysis and improves metabolic control in diet-induced obesity. bioRxiv, 2025.2005.2009.653092 (2025).

12. E. Staal, C. Faurholt, Studies on carbamates. IV. The carbamates of hydrazine. Dan. Tidsskr. Farm. 25, 1–11 (1951).

13. A. El-Faham, F. Albericio, Peptide Coupling Reagents, More than a Letter Soup. Chem. Rev. 111, 6557–6602 (2011).

14. C. Gibson, S. L. Goodman, D. Hahn, G. Hölzemann, H. Kessler, Novel Solid-Phase Synthesis of Azapeptides and Azapeptoides via Fmoc-Strategy and Its Application in the Synthesis of RGD-Mimetics. J. Org. Chem. 64, 7388–7394 (1999).

15. Y. Garcia-Ramos, W. D. Lubell, Synthesis and alkylation of aza-glycinyl dipeptide building blocks. J. Pept. Sci. 19, 725–729 (2013).

16. K. Tarchoun, M. a. Yousef, Z. Bánóczi, Azapeptides as an Efficient Tool to Improve the Activity of Biologically Effective Peptides. Future Pharmacol. 2, 293–305 (2022).

17. M. A. McMechen, E. L. Willis, P. C. Gourville, C. Proulx, Aza-Amino Acids Disrupt β-Sheet Secondary Structures. Molecules 24, 1919 (2019).

18. S. Pechenov et al., Development of an orally delivered GLP-1 receptor agonist through peptide engineering and drug delivery to treat chronic disease. Sci. Rep. 11, 22521 (2021).

19. L. B. Knudsen, J. Lau, The Discovery and Development of Liraglutide and Semaglutide. Front. Endocrinol. (Lausanne) 10 (2019).

20. T. C. Dinsmore et al., Potent and Protease Resistant Azapeptide Agonists of the GLP-1 and GIP Receptors. Angew. Chem. Int. Ed. 63, e202410237 (2024).

21. Y. Al-Abed, K. F. Cheng (2020) Preparation of O-benzotriazole and O-imidazole synthons for use in the synthesis of peptidomimetics including azapeptides. (The Feinstein Institutes for Medical Research).

